# Developmental pyrethroid exposure in mouse leads to disrupted brain metabolism in adulthood

**DOI:** 10.1101/2023.10.13.562226

**Authors:** Melissa A. Curtis, Nilanjana Saferin, Jennifer H. Nguyen, Ali S. Imami, William G. Ryan, Kari L. Neifer, Gary W. Miller, James P. Burkett

**Affiliations:** Department of Neurosciences, University of Toledo College of Medicine and Life Sciences, Toledo, OH 43614; Department of Environmental Health, Emory Rollins School of Public Health, Atlanta, GA 30322; Department of Environmental Health Sciences, Mailman School of Public Health, Columbia University, New York, NY 10032 (current)

**Keywords:** pyrethroids, developmental exposure, metabolomics, brain, pyrethroids, developmental exposure, metabolomics, brain

## Abstract

Environmental and genetic risk factors, and their interactions, contribute significantly to the etiology of neurodevelopmental disorders (NDDs). Recent epidemiology studies have implicated pyrethroid pesticides as an environmental risk factor for autism and developmental delay. Our previous research showed that low-dose developmental exposure to the pyrethroid pesticide deltamethrin in mice caused male-biased changes in the brain and in NDD-relevant behaviors in adulthood. Here, we used a metabolomics approach to determine the broadest possible set of metabolic changes in the adult male mouse brain caused by low-dose pyrethroid exposure during development. Using a litter-based design, we exposed mouse dams during pregnancy and lactation to deltamethrin (3 mg/kg or vehicle every 3 days) at a concentration well below the EPA-determined benchmark dose used for regulatory guidance. We raised male offspring to adulthood and collected whole brain samples for untargeted high-resolution metabolomics analysis. Developmentally exposed mice had disruptions in 116 metabolites which clustered into pathways for folate biosynthesis, retinol metabolism, and tryptophan metabolism. As a cross-validation, we integrated metabolomics and transcriptomics data from the same samples, which confirmed previous findings of altered dopamine signaling. These results suggest that pyrethroid exposure during development leads to disruptions in metabolism in the adult brain, which may inform both prevention and therapeutic strategies.

**Highlights:**

- Developmental pyrethroid exposure disrupts brain metabolism in adulthood
- Exposure disrupts metabolic pathways for folate, retinol, and tryptophan
- Exposure disrupts genetic and metabolic pathways for dopamine signaling

## 1. Introduction

Neurodevelopmental disorders (NDDs) are a cluster of lifelong brain disorders including autism spectrum disorders, attention deficit hyperactivity disorder (ADHD), learning disabilities, sensory impairments, schizophrenia, and intellectual disabilities^1,2^. These are often incurable disorders with limited biomarkers and treatment options^3^. The prevalence of NDDs is rapidly rising, affecting as many as 17% of children in the United States^4^. Research has historically focused on genetic factors in the etiology of NDDs, however, recent studies have shown that genetic anomalies can only account for 30-40% of all cases, and that environmental risk and gene-environment interactions may contribute substantially the risk for autism^5–9^. Environmental toxicants such as air pollution^10^, heavy metals (lead and mercury)^11^, persistent organic pollutants (DDT and polychlorinated biphenyls)^12,13^, flame retardants (polybrominated diphenyl ethers and organophosphate esters)^14,15^, and pesticides (organophosphates and pyrethroids)^16^ have been implicated in risk for NDDs in humans^17,18^. Understanding the risk factors associated with NDDs and their underlying molecular mechanisms is critical for prevention and the development of therapeutic strategies.

One potential environmental factor of concern is developmental exposure to pyrethroid pesticides. Pyrethroids are neurotoxic chemicals commonly used for agriculture and mosquito fogging, and in household applications such as insect sprays, flea collars, and head lice shampoos^19^. Pyrethroids are thought to be safe for adults and have low environmental impact, making them one of the most widely used pesticide classes in the US, with 70-80% of the general US population having detectable pyrethroid metabolites in their blood^20,21^. Nonetheless, epidemiology studies have shown that prenatal exposure to pyrethroids is a risk factor for NDDs^16,22,23^ and that pyrethroid metabolite levels in the blood of pregnant women is associated with parent-reported mental development in the child^24^. Moreover, pyrethroid metabolites in blood and urine are associated with the risk for ADHD in children^20,21^.

Mouse models have recently been developed to study comparable effects of developmental exposure to deltamethrin, a type II pyrethroid, in humans^20,25–27^. These models reproduce the oral route of exposure, which is the most common route of chronic exposure in humans; and, to control for bioaccumulation, exposures are performed every 3 days, an interval at which the pesticide and its metabolites are expected to be entirely cleared. Deltamethrin exposure in mice during pregnancy and lactation at doses well below the Environmental Protection Agency (EPA) benchmark dose of 14.5 mg/kg is sufficient to cause a broad range of changes in the brain and behavior of the offspring. Behaviorally, deltamethrin exposure during development results in hyperactivity, repetitive behaviors, reduced vocalizations, impulsivity, and learning deficits^20,25^. In the brain, developmental deltamethrin exposure leads to broad changes in striatal dopamine signaling, altered levels of voltage-gated sodium channels, elevated endoplasmic reticulum stress, altered brain-derived neurotrophic factor (BDNF) levels, and changes in molecular pathways for circadian rhythms and synaptic plasticity^26,28–33^. Behavioral effects in this model primarily impact male mice, and molecular effects are similarly male-biased. However, the underlying processes altered by deltamethrin exposure to drive these broad changes are unknown, hindering the identification of effective therapeutic targets or the identification of biomarkers.

Metabolomics – the quantitative measure of all low-molecular-weight biological molecules detectable in a sample using advanced analytical chemistry techniques^34^ – is beginning to be recognized as an invaluable tool for translational research related to biomarker discovery, drug target discovery, drug responses, and disease mechanisms^35^. Mass spectrometry-based metabolomics detects and quantifies thousands of molecular features simultaneously, which are identified by a combination of retention time and signature^36^. These molecular features are then mapped onto specific metabolites using annotated reference databases. Metabolomics has the potential to identify metabolic processes that are disrupted by developmental exposure to pesticides in an unbiased manner, providing the foundation for testable hypotheses identifying biomarkers or drug targets and describing mechanisms driving the broad changes in the brain observed with developmental pesticide exposure.

In this study, we exposed pregnant mouse dams to a low level of deltamethrin (3 mg/kg every 3 days) throughout pregnancy and lactation. This dose was chosen based on prior studies showing behavioral and neurological effects in offspring at this developmental dose^20,25,33^ and because it is substantially lower than both the EPA-determined benchmark dose^19^ and the developmental no observable adverse effect level^37^. We chose to focus on male offspring because both the behavioral and molecular outcomes of developmental pyrethroid exposure (DPE) have been male-biased, so the most significant changes in brain metabolism would be expected to occur in males. Deltamethrin readily penetrates all regions of the brain following acute oral exposure in mice, potentially affecting many brain regions^38^; and following acute oral exposure in pregnant mice, the related pyrethroid permethrin crosses the placenta into the embryo, where it potentially accesses the entire developing brain^39^. We therefore selected the whole brain as our level of investigation in order to best represent major changes in brain metabolism across many brain regions and majority cell types. In line with these decision points, we performed untargeted high-resolution metabolomics on whole-brain tissue collected from the adult male offspring. As a cross-validation method, we performed transcriptomics on split brain samples from the same subjects. We hypothesized that low-dose developmental pyrethroid exposure would lead to changes in metabolic processes in the adult brain.

## 2. Materials and Methods

### 2.1. Animals

Experimental subjects were the adult (P56+) male offspring of female C57BL6/J mouse dams. Virgin dams were kept on a 12/12 hour light cycle and given access to water and high-fat breeder food ad libitum. Dams were housed in female pairs, then in trios with a male sire until 3 days prior to birth, and finally singly housed through birth and weaning. Offspring were weaned at 20-22 days of age into single-sex cages of 3-5 mice and were socially housed continuously into adulthood. All procedures were approved by the Emory University IACUC and performed in accordance with the US Animal Welfare Act.

### 2.2. Chemicals

Deltamethrin (Sigma, St. Louis, MO) is the reference pyrethroid selected by the EPA (EPA, 2011). Aliquots were prepared by dissolving deltamethrin (Sigma, St. Louis, MO) in acetone (Sigma), mixing with corn oil (Mazola, Memphis, TN), allowing the acetone to evaporate overnight, and then storing at -80° C in capped borosilicate glass vials (Fisher Scientific, Hampton, NH) until the day of use. Aliquots of vehicle were prepared by allowing a similar volume of acetone to evaporate in corn oil.

### 2.3. Study design

The study design consisted of two independent groups: pesticide exposed and vehicle control. Mouse dams were exposed to deltamethrin (or vehicle) during preconception, pregnancy, and lactation as described below. The experimental groups for these experiments were composed of a single adult male offspring from each of the resulting litters.

### 2.4. Experimental units

Since experimental exposure was given to the dam, the experimental unit was the dam/litter, and each dam/litter was counted as N=1. A single male offspring was chosen at random to represent the litter.

### 2.5. Elimination criteria

Dams and their litters were eliminated from the study if they did not consume 90% of the peanut butter containing the pesticide or vehicle during 90% of the feedings (see below), if they did not give birth within 28 days of being paired with a sire, or if their litter did not contain at least one male and one female pup.

### 2.6. Sample size

A total of 40 female mice, pre-screened for peanut butter preference, (N=20 per group) were offered experimental or control treatment in this study. Three dams were eliminated due to inconsistent consumption, and two dams failed to produce a litter within 28 days of pairing. Four whole-brain samples were eliminated for litter size/composition. This left N=18 exposed and N=13 control samples.

### 2.7. Developmental pyrethroid exposure (DPE)

Mice were developmentally exposed to the pyrethroid pesticide deltamethrin as previously described^25^. Starting two weeks prior to being paired with a male, dams were fed deltamethrin (3 mg/kg by voluntary oral consumption in peanut butter/corn oil) or vehicle once every 3 days, continuing throughout pre-conception, pregnancy, and weaning. Dams were weighed immediately prior to each feeding, and the appropriate amount of corn oil was mixed with JIF peanut butter (J.M. Smucker, Orville, OH) to achieve the desired dose. To ensure that cagemates and offspring did not consume the peanut butter, adult cagemates were temporarily separated and peanut butter was provided in an elevated position. Dams that failed to consume 90% of the peanut butter during 90% of the feedings were eliminated from the study. Offspring were only exposed to deltamethrin indirectly through the dam. Offspring were weaned from the dam as above and received no additional treatment until experimental testing.

### 2.8. Tissue

Adult male offspring were sacrificed by rapid decapitation and dissected. The brains were flash-frozen using liquid nitrogen and pulverized. The resulting powder was aliquoted for use in split-sample multiomics^33^.

### 2.9. Untargeted high-resolution metabolomics

Pulverized brain tissue aliquots were shipped on dry ice to the HERCULES High Resolution Metabolomics Core at Emory University. Sample preparation and high-resolution tandem liquid chromatography-mass spectrometry (LC-MS) were performed by the Core. The Metabolomics Core sent the results of LC-MS metabolomics in the form of raw feature tables for both c18neg and hilicpos modes, which were generated by the Core using xMSanalyzer and xMSannotator^40,41^. As this was an untargeted analysis, the Metabolomics Core included quantitative standards only for a small subset of common metabolites. Therefore, we considered the pathway analysis to be the primary outcome.

#### 2.9.1. Metabolomics preprocessing

The R package xmsPANDA (version 1.3.2) was used for data filtering of features with missing values, log2 transformation, and quantile normalization^42^. xmsPANDA was then used to identify features of interest based on two criteria: (1) the feature is significantly different between the two groups (p<0.05 uncorrected, Limma t-test); and (2) the feature is important in discerning between the two groups, as determined by a variable importance in the projection (VIP) score >2 (partial least-squares discriminant analysis). VIP scores, p-values and log fold change were used to generate volcano and Manhattan plots and to conduct unsupervised two-dimensional hierarchical clustering analysis.

#### 2.9.2. Metabolite annotation

Features of interest were mapped to the KEGG and HMDB databases using the web-based tool, MetaboAnalyst 5.0^43,44^. Feature tables from xmsPANDA were used as input into the tool’s modules for data filtering, normalization, feature selection, and statistical analysis. Mummichog pathway analysis was used with default parameters as the primary outcome of interest.

#### 2.9.3. Metabolite abundance

Because we chose an untargeted metabolomics approach, the Metabolomics Core included quantitative metabolite standards only for a small list of common metabolites. Therefore, we considered metabolite abundance as a descriptive analysis. The xmsPANDA (version 1.3.2) “quant” function was used to calculate individual metabolite abundances (in µM) using these pre-quantified metabolite standards. Raw feature tables (including the standards) were input into the quant function with a mass tolerance of 5 ppm, a retention time of 30 seconds, and all other parameters set to default.

### 2.10. Transcriptomics data analysis

Previously existing data from split-sample transcriptomics on the same brain samples was used to complement and cross-validate the metabolomics data^33^. Briefly, aliquots of the same frozen brain samples used in metabolomics were homogenized, the lysate centrifuged and filtered, and aliquots prepared using the RNEasy Plus Micro Kit (QIAGEN, Germantown, MD). Extracted RNA aliquots were sent to the University of Michigan Advanced Genomics Core for quality control, library preparation, and next generation sequencing. The University of Michigan provided paired end FASTQ files, which we aligned to the GRCm38 mouse genome using Hisat2^45^, counted using GenomicAlignments and GenomicFeatures^46^, and compared between groups using Deseq2^47^. A variance partition^48^ was used to select genes of interest that were both differentially expressed between groups (p<0.05 uncorrected) and had at least 15% of the variance in expression explained by group.

### 2.11. Transcriptome Perturbagen Analysis

Post-processed differential gene expression data were compared to the iLINCS online database using concordant and discordant perturbagen analysis^49,50^. The R packages drugfindR (version 0.99.556) and biomaRt (version 2.60) were used. The following parameters were used: Output_lib =“CP”, filter threshold =0.075, similarity threshold = 0.321. P values were converted to FDR-corrected q values using the total database size of 143,374 chemical perturbagen signatures.

### 2.12. Integration of transcriptome and metabolome data sets

Split-sample transcriptome and metabolome data from the same subjects were integrated to provide a cross-validated set of differentially represented pathway clusters. We used the MetaboAnalyst Joint Pathway Analysis^43^ to combine our list of genes of interest with our list of annotated metabolites of interest. The result is a list of differentially represented pathways with q-values, or false discovery rate corrected p-value. Differentially expressed pathways (q-value < 0.05) were clustered and visualized using PAVER^51^ (version 0.0.0.9000). PAVER generated UMAP scatter plots of clustered pathways and derived names for the clusters by selecting the term most similar to each cluster’s average embedding. Heatmaps were generated showing clusters organized by average cluster enrichment, with pathways inside the cluster organized by the enrichment score, which is equal to -log_10_ of the q-value (the FDR-corrected p-value).

## 3. Results

### 3.1. Feature analysis

We performed untargeted high-resolution metabolomics on whole-brain tissue from adult male DPE mice. Mass spectrometry detected 120,416 unique molecular features in the positive and negative ion modes (Supplemental File 1A-1B). Preprocessing of the raw data identified 731 molecular features of interest that were significantly different between DPE mice and controls (p<0.05 uncorrected) and were important in identifying differences between the groups (VIP > 2) (Figure 1A-B; Supplemental Figures S1-S3). Hierarchical clustering analysis using these 731 features was able to confidently separate the two groups (adjusted r=0.37; p=0.040; sample average silhouette width=0.12; Figure 1C). These 731 features mapped on to 116 annotated metabolites of interest (Supplemental File 1C). These results indicate that developmental pyrethroid exposure causes a change in molecular features of interest relative to controls.

**Figure 1.**
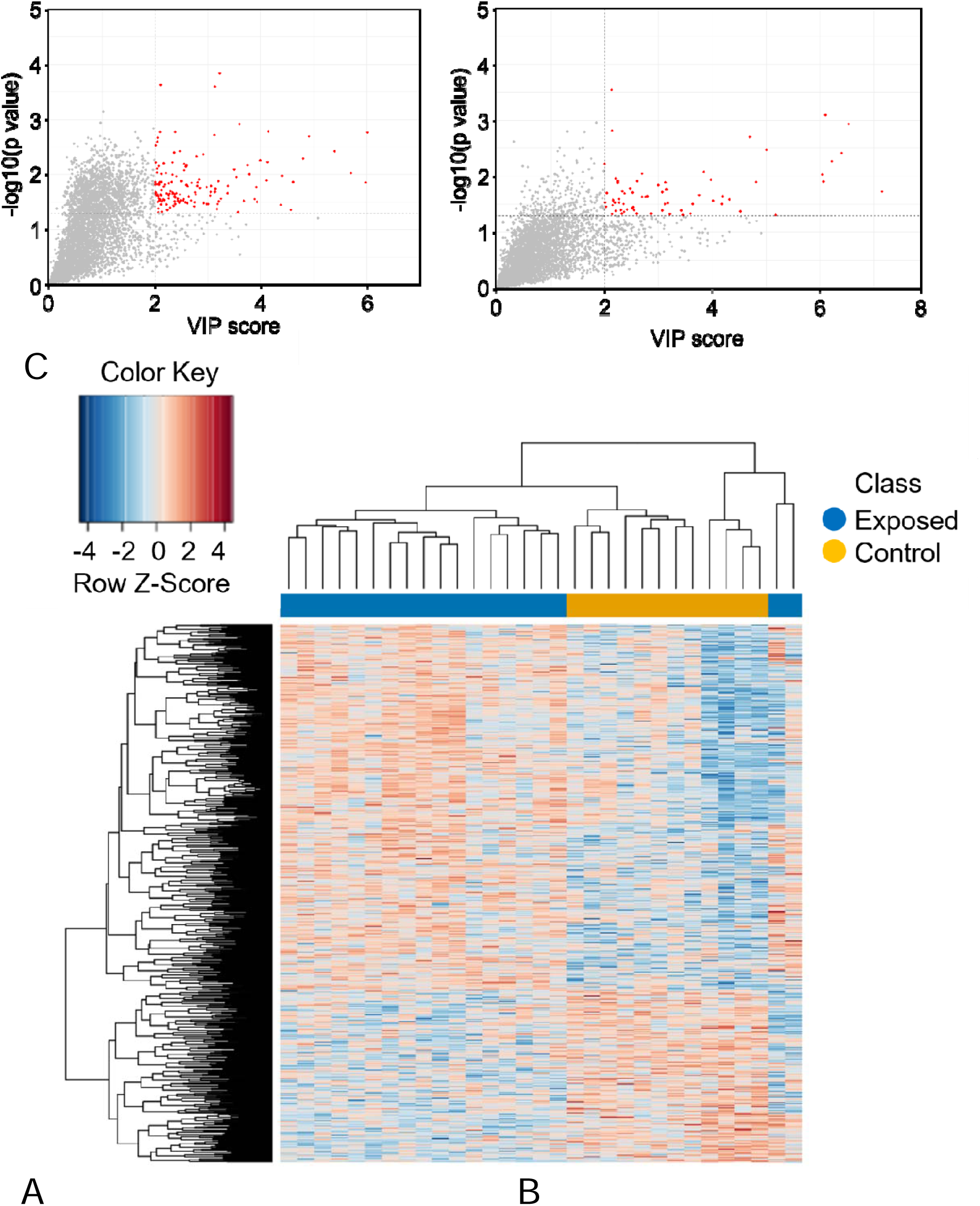
Molecular features of interest. Preprocessing of raw feature tables identified (A) positively charged and (B) negatively charged molecular features that were significantly different between groups (by p-value) and important in modeling differences between the groups (by VIP score). (C) A heat map shows the hierarchical clustering analysis that was performed using features of interest, which was able to confidently separate the two groups (adjusted r=0.37; p=0.040). Group subjects are clustered along the x-axis, with features of interest (mz/rt) on the y-axis.

**Figure 2.**
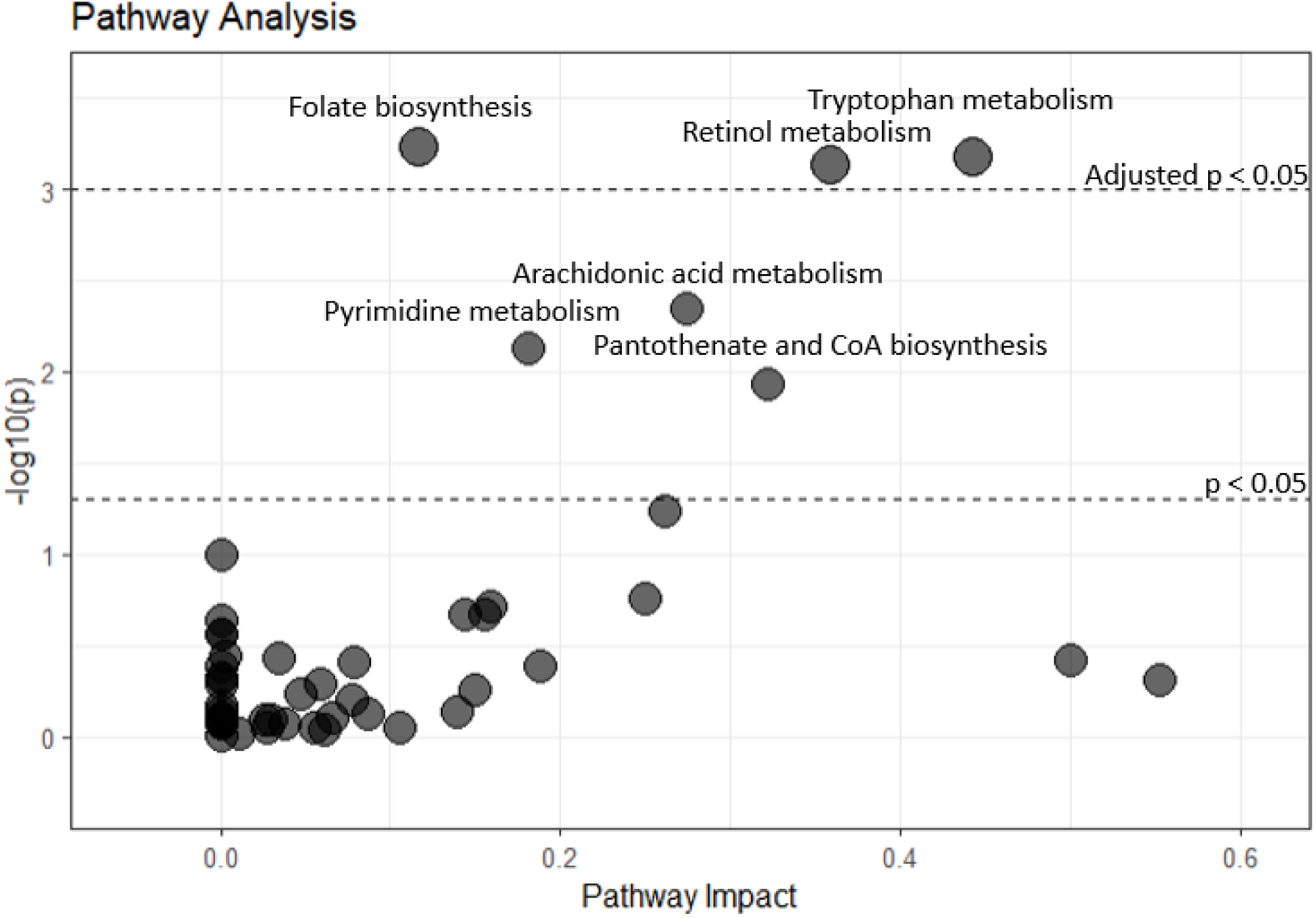
Metabolite pathway analysis. Pathway analysis on the 116 annotated metabolites of interest was used to map the metabolites to metabolic pathways. Pathways were evaluated by statistical significance (p-value) and the enrichment of metabolites in the pathway (pathway impact). Dashed horizontal lines show the cutoff values for raw p < 0.05, and p < 0.05 after adjustment for false discovery rate.

### 3.2. Metabolite Pathway Analysis

Pathway analysis was considered the primary outcome of interest for metabolomics due to the lack of broad quantitative standards in the untargeted approach. The 116 annotated metabolites of interest were used to identify metabolic pathways that were significantly altered in DPE mouse brain. Metabolites for folate biosynthesis, retinol metabolism, and tryptophan metabolism were significantly different between DPE mice and controls (hypergeometric test, p<0.02; Figure 3, Supplemental Figures S4-S6, Supplemental File 1D). Separately, a descriptive abundance analysis on the 94 metabolites for which quantitative standards were included was performed, only four of which mapped on to the pathways of interest (Figure 3, Supplemental File 1E-F). Retinoate (i.e. retinoic acid) was decreased in relative abundance in DPE mouse brain (DPE: 5.4 ± 0.7 uM, Control: 7.7 ± 0.4 uM; t-test, t=2.85, p=0.0079 uncorrected; Figure 3), suggesting that the overall direction of change in retinol metabolism may have been downward.

**Figure 3.**
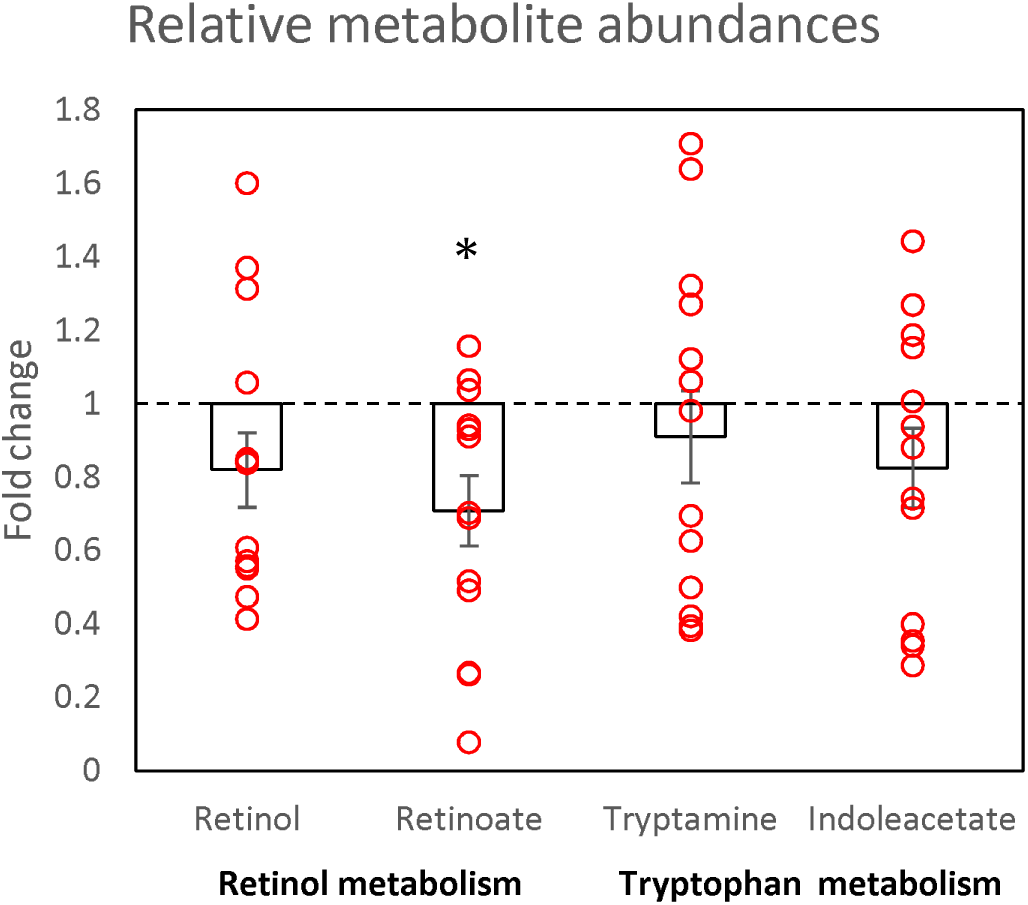
Descriptive analysis of relative abundance (by fold change from control) of metabolites of interest found in retinol metabolism and tryptophan metabolism pathways. Two metabolites were identified in each of these pathways by abundance analysis. Retinoate (i.e. retinoic acid) was reduced by 30% in DPE mouse brain relative to controls. Error bars show standard error. * p < 0.05 vs. control.

### 3.3. Multiomics integration of transcriptome and metabolome

As a form of cross-validation, we used split-sample transcriptome and metabolome data to generate a set of multi-modally dysregulated pathway clusters. Transcriptome data were previously published and analyzed^27^. The variance partition method identified 65 genes of interest that were differentially expressed between groups and had >15% of the variance explained by group (Supplemental File 1G). Multimodal integration of these 65 genes of interest with the 116 annotated metabolites of interest identified 184 over-represented KEGG pathways (Supplemental File 1H). These pathways segregated into seven pathway clusters, including dopaminergic synapse and MAP kinase signaling, among others (Figure 4A-B).

**Figure 4.**
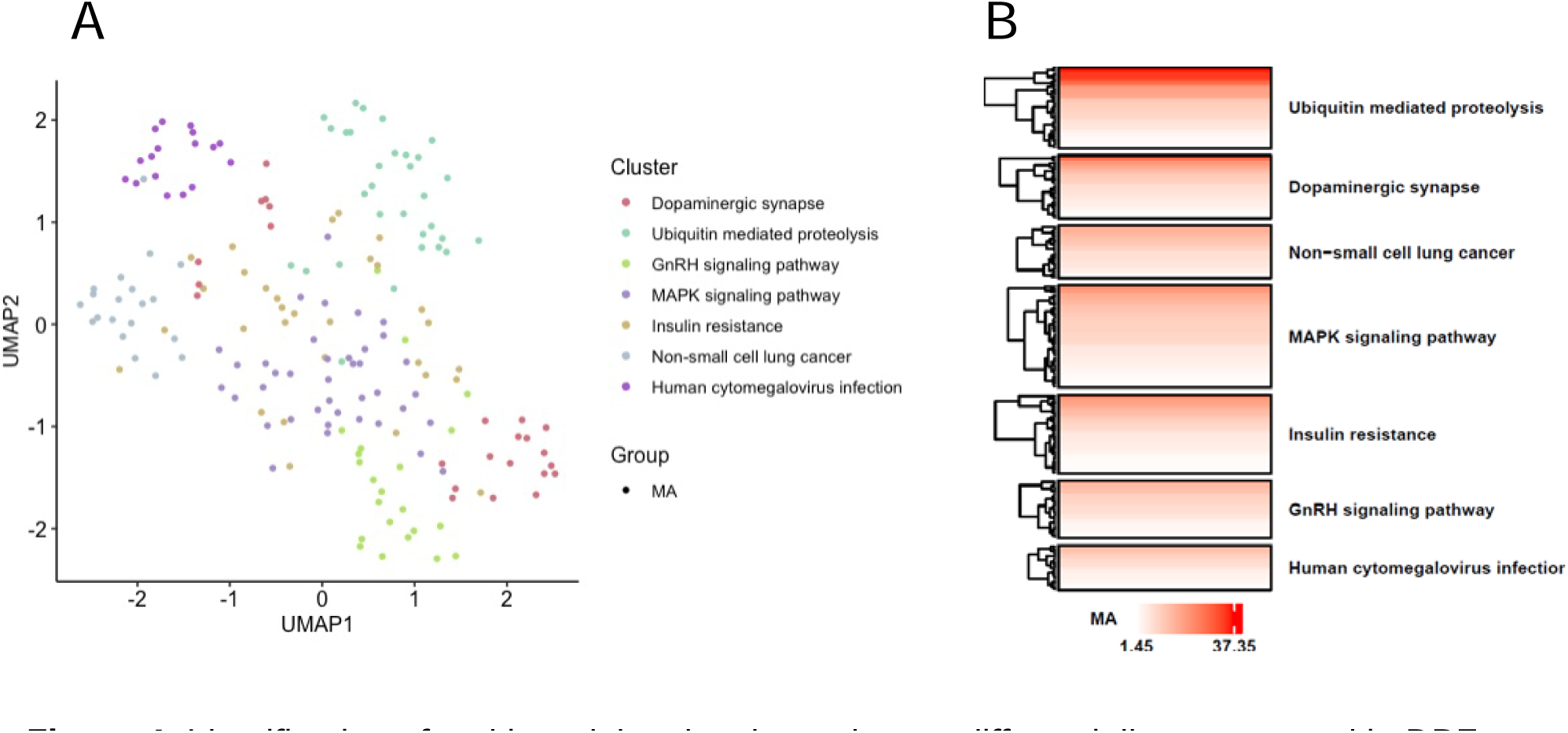
Identification of multi-modal molecular pathways differentially represented in DPE mouse brain using multiomics integration. (A) UMAP components analysis showing seven clusters of molecular pathways dysregulated in DPE mouse brain. (B) A pathway cluster heatmap shows the clusters internally sorted by enrichment score (-log_10_ of q-value) and arranged from most enriched (top) to least enriched (bottom).

### 3.4. Perturbagen Analysis

We compared the split-sample transcriptome data from DPE mice to the iLINCS online database of compound-induced transcriptional changes using perturbagen analysis (Table 1A-B, Supplemental File 1I). Among the primary “concordant” perturbagens (having a similar transcriptional signature as DPE) were trimethoprim (similarity = 0.5459), a folate synthesis inhibitor^52^; and AG14361 (similarity = 0.5073), an inhibitor of mammalian target of rapamycin (mTOR) signaling and apoptosis^53,54^ (Table 1A). Among the primary “discordant” perturbagens (having the opposite transcriptional signature as DPE) were pelitinib (similarity = - 0.5473), an epidermal growth factor receptor (EGFR) inhibitor^55^; and Brefeldin A (similarity = - 0.4512), an inhibitor of vesicle formation and synaptic vesicle recycling^56,57^ (Table 1B).

**Table 1.** Concordant and discordant perturbagens. Perturbagen analysis showed the similarity between the transcriptome profiles of DPE mouse brain tissue (compared to control) and “perturbagen” drugs applied to cell culture (compared to control) from the iLINCS database. Similarity scores are Pearson r values, with larger positive or negative values indicating more correlation between data sets. Significance is reported as FDR-corrected q values. (A) Concordant perturbagens with a positively correlated transcriptome profile. (B) Discordant perturbagens with a negatively correlated transcriptome profile.

| **Concordant Perturbagen** | **Similarity** | **FDR** |
| --- | --- | --- |
| *Trimethoprim* | 0.5459 | $2.09 \times 10^{-8}$ |
| *Erlotinib* | 0.5145 | $4.89 \times 10^{-7}$ |
| *AG 14361* | 0.5073 | $7.13 \times 10^{-7}$ |
| *Levosalbutamol* | 0.4975 | $1.44 \times 10^{-6}$ |
| *Betahistine* | 0.4889 | $2.95 \times 10^{-6}$ |

**Table 1.** Concordant and discordant perturbagens.
| **Discordant Perturbagen** | **Similarity** | **FDR** |
| --- | --- | --- |
| *Pelitinib* | -0.5473 | $2.09 \times 10^{-8}$ |
| *CHEMBL2143628* | -0.5061 | $7.13 \times 10^{-7}$ |
| *Brefeldin A* | -0.4512 | $3.48 \times 10^{-5}$ |
| *IM-12* | -0.4398 | $5.96 \times 10^{-5}$ |
| *Sorafenib* | -0.4423 | $7.55 \times 10^{-5}$ |

## 4. Discussion

Adult mice exposed during development to low levels of deltamethrin showed a pattern of altered small molecule metabolites in three key metabolic pathways in whole brain. Compared to vehicle-exposed controls, DPE mice showed altered folate biosynthesis, a process that is particularly critical during neurodevelopment for preventing neural tube defects. Additionally, the whole-brain transcriptional profile of adult DPE mice showed a high similarity to the transcriptional profile associated with trimethoprim, a drug that inhibits folate synthesis. These results suggest a multi-modal effect of pyrethroid exposure during development on subsequent adult folate metabolism at the whole-brain level. DPE mice also had altered retinol and tryptophan metabolism, two processes related to circadian rhythms, in line with other research showing reduced whole-brain gene transcription of clock genes in DPE mice^33^. DPE mouse brain had reduced retinoic acid concentration as compared to controls, suggesting that retinoic acid signaling and metabolism may be decreased. Additionally, multiomics integration of metabolome and transcriptome revealed altered molecular pathways for dopaminergic synapses, validating these findings against prior literature on dopamine dysfunction in DPE mice^20,25^. This is the first study using an unbiased metabolomics approach to identify effects of DPE on the brain in need of further investigation in future hypothesis-driven studies.

Although specific exposure levels in blood and tissues of pregnant mice were not measured, other studies provide context. For example, in adult rats, a 1 mg/kg dose of deltamethrin resulted in a serum concentration of 56 ng/mL^58^, with human models predicting double these levels^59^. Comparatively, pregnant mice exposed to 1.5 mg/kg permethrin had serum concentrations of 260 ng/mL^39^. Deltamethrin levels in the general population in Beijing ranged from 0 to 17.34 ng/mL^60^, while pregnant Chinese women had an average level of 39 ng/mL^61^. In the US, urinary metabolite levels increased from 0.292 ng/mL in 1999–2000^62^ to 660 ng/mL in 2011–2012^63^. These statistics suggest that our chosen exposure dose may represent an above-average exposure, but one which importantly falls below the currently established developmental no observable adverse effect level^37^ and the EPA-set benchmark dose^19^.

Folate is critical for early neural development and is universally recognized as a key prenatal vitamin for the prevention of neural tube defects. Prenatal folate supplementation in humans decreases the risk for autism, developmental delay, and other NDDs^64,65^, and mouse models of maternal folate deficiency consistently show increases in neural tube defects and tumor risk^66,67^. Additionally, high-dose prenatal folate supplementation in humans may be protective against the increased risk for NDDs contributed by pyrethroid pesticide exposure^68^. In our study, male mice exposed to deltamethrin during development showed altered folate metabolism in the brain in adulthood. Trimethoprim, a folate synthesis inhibitor, was also the most significant hit in our concordant perturbagen analysis, which compared transcriptomic profiles of gene expression between the iLINCS database of drug effects on cell culture, and split samples of whole brain from our mouse subjects. The effect of DPE on adult folate metabolism and folate-relevant transcriptomic profile in the mouse brain may represent a primary cause of the effects of DPE on NDD-relevant behaviors^25^ and on NDD-relevant molecular pathways for synaptic plasticity^33^. Folate is also a precursor in dopamine biosynthesis, which may explain the altered multiomic pathways for dopaminergic synapses, as well as previous findings of broad changes in the dopamine system in DPE mice^20,25^.

Perturbagen analysis also identified Brefeldin A, an inhibitor of vesicle formation and synaptic release, as having a highly dissimilar transcriptome profile to that of the DPE mouse brain. Multiomics analysis also showed alterations in metabolic and transcriptional pathways for “dopamine synapse,” and previous findings have highlighted hyperactive dopamine signaling in the striatum of DPE mice^20,25^. Therefore, this discordant profile may reflect a broad increase in synaptic dopamine release in DPE mouse brain. The most highly dissimilar transcriptomic profile to DPE was Pelitinib, an EGFR inhibitor; potentially suggesting a relative increase in growth factor signaling in DPE mouse brains. This would synergize with the relative similarity of the transcriptomic profile of AG14361, an inhibitor of apoptosis-related mTOR signaling. However, the presence of an EGFR inhibitor (Erlotinib) on the concordant perturbagen list as well makes this finding difficult to interpret.

Tryptophan metabolism was also altered in the brains of adult DPE mice. Tryptophan is an essential amino acid and is metabolized along two metabolic pathways: a small amount of tryptophan is metabolized via the methoxyindole pathway, which produces serotonin and melatonin; while the majority of tryptophan is metabolized via the kynurenine pathway, which regulates immune response and is considered neuroprotective^69,70^. Placental tryptophan metabolism is an essential source of serotonin in fetal neurodevelopment^71–73^, and disruptions in serotonin signaling during fetal brain development cause a wide range of neural defects^74^. The kynurenine pathway generates two compounds, kynurenic acid and quinolinic acid, and disruptions in both of these compounds have been associated with NDDs and autism^75^. Furthermore, acute deltamethrin exposure in rats decreases kynurenic acid production by 31%, suggesting that deltamethrin may have a direct effect on this pathway^76^.

Both altered retinol metabolism and altered melatonin (via tryptophan metabolism) in DPE mouse brains may provide links between DPE and altered circadian rhythms. DPE mice show reduced transcription of critical clock genes at the whole-brain level^33^. During development, retinoids play a key role in neuronal differentiation, neural tube development, and neurite outgrowth^77^. In adulthood, retinol metabolism and retinoic acid are critical for the regulation of both circadian and seasonal rhythms^78^. Melatonin also plays an essential role in circadian rhythms in the brain and is one of the most reliable markers of the sleep-wake cycle^79^.

Disruption of circadian rhythms and sleep are among the most common comorbidities in developmental disorders^80,81^. Retinoic acid is also highly expressed in the dopamine system and is a key regulator of dopamine synthesis and signaling^82,83^, both of which are disrupted in DPE mice^20,25^. Finally, retinoic acid is also critical to adult neurogenesis and synaptic plasticity^84–86^, and alterations related to these two processes were detected by both kinomic and transcriptomic analysis of these same brains^27^. Thus, our non-biased metabolomics study adds to the literature highlighting retinol and tryptophan metabolism as worthy targets for future investigations into the long-term effects of developmental pyrethroid exposure.

Multiomics integration of metabolome and transcriptome served to validate and reinforce these findings against prior literature. The most enriched cluster of molecular pathways in the integrated data set was ubiquitin-mediated proteolysis, a critical mechanism of autophagy and apoptosis. Perturbagen analysis of the transcriptome also found the mTOR inhibitor AG14361 to have a similar transcriptional profile to DPE, suggesting a decrease in apoptosis-related transcription. Apoptotic pathways were also prominent in the kinomics analysis from these same subjects^33^, where it appeared that cellular growth was upregulated and apoptosis downregulated. Changes in the mitogen-associated protein kinase (MAPK) signaling pathway were also detected in kinomics analysis, where it appeared that both extracellular signaling kinase (ERK) and rapidly accelerated fibrosarcoma (RAF) kinase activity was upregulated by DPE. The dopaminergic synapse cluster of pathways also validates these findings against prior literature showing alterations of dopamine, dopamine receptors, dopamine transporter, and dopamine release dynamics in low-dose DPE in mice^20,25^.

There are key strengths and limitations to this study that must be considered. A significant limitation of this study is its use of a single exposure dose of deltamethrin, selected based on previous research and significantly below both regulatory benchmark doses and developmental no observable adverse effect levels^20,25,33^. A careful look at dose and the dose-dependent effects of DPE is a critical next step. Another limitation of this study is the use of whole-brain tissue as the level of analysis. We chose to analyze whole brain because pyrethroid pesticides readily penetrate all regions of the brain and cross the placenta into the embryo following acute oral exposure in mice^38,87^. Therefore, examining whole-brain tissue strengthens our ability to see metabolic effects permeating every brain region and dominant cell type, which are also the most likely to be good targets for oral pharmacological treatment. However, more specific effects on individual brain regions or minority cell types are unlikely to be represented in our data and are an important future direction. Based on the DPE phenotype in mouse of hyperactivity, repetitive behaviors, and cognitive deficits, a primary effect on prefrontal cortex, anterior cingulate cortex, and striatum might be predicted; and effects on circadian rhythm molecular pathways directly implicate the suprachiasmatic nucleus. Finally, our study looked at the effects of DPE exclusively on male mouse brains. This decision was primarily driven by the male-biased effects of DPE in prior studies in mice^20,25^, suggesting that the largest changes would be expected in male mouse brains. However, as a consequence, this study cannot contribute to the larger literature on male-biased sensitivity to neurotoxicants because it lacks a female comparison. Including female subjects in future studies would allow us to see sex-dependent changes in brain metabolism and further identify the root causes of the male bias in the changes in NDD-relevant behaviors in DPE mice.

### 4.1. Conclusions

Pyrethroid pesticides are among the most commonly used in the US, and the exposure level of the general public is high. If developmental exposure to pyrethroid pesticides increases risk for autism and other developmental disorders, this represents a critical public health challenge. Here, we used an unbiased approach to show that developmental exposure to a low level of deltamethrin in mice caused changes in brain metabolism related to folate, tryptophan, and retinol metabolism, which are strongly related to developmental disorders in humans. We also showed that the transcriptomic profile resulting from developmental pyrethroid exposure resembles the action of folate-related drugs; and that multiomics integration of these two datasets identifies molecular pathways for dopamine signaling, proteolysis, and MAPK signaling. Much work remains to verify these findings using specific targeted methods, which may point to potential avenues for prevention and treatment.

## Acknowledgements

We gratefully acknowledge the Emory University HERCULES High Resolution Metabolomics Core, and in particular Dr. Dean Jones and Sami Teeny, for their significant assistance with data analysis. We would also like to thank Dr. Sarah Daniel of In Scripto, LLC for comments on this manuscript.

## Authorship contribution statement

JPB and GWM designed the exposure study, and JPB designed the analysis strategy. All samples were collected in the lab of GWM by JPB and transferred to the lab of JPB. MAC processed all samples, distributed them to the core facilities, and coordinated the initial analysis of all data. JHN and ASI participated in the final analysis of all data and assembly of figures under the guidance of JPB. JPB, MAC, and JHN interpreted all results. NS performed background research and prepared the manuscript. KLN provided experimental, administrative, and managerial support. All authors participated in writing the manuscript, approved the final version, and share responsibility for the content. No generative AI or AI-assisted technologies were used in the writing of this manuscript.

## Data statement

All raw data for metabolomics, including all metadata, are available in the supplementary materials. All sequencing data are available on NCBI GEO (GSE241185).

