## Supplementary Figures for "Developmental pyrethroid exposure in mouse leads to disrupted brain metabolism in adulthood"

**This PDF file includes:**

Figures S1 to S6
Table S1

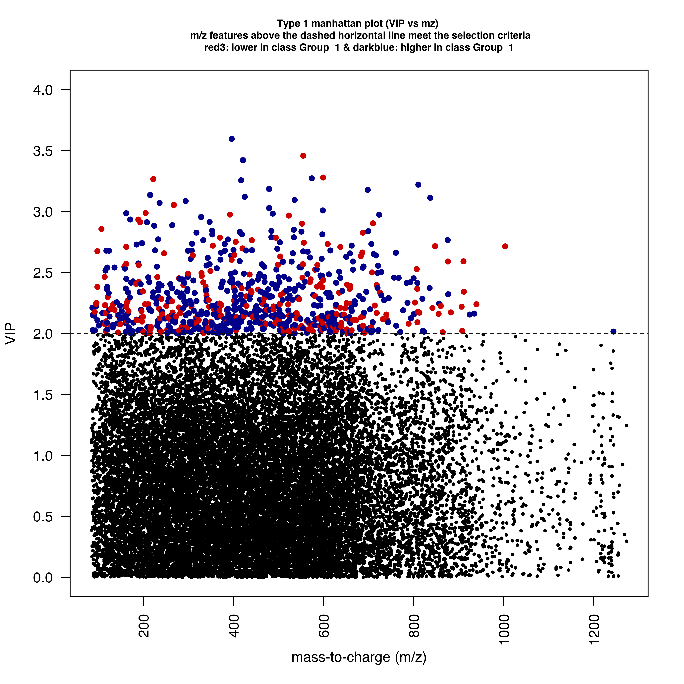

Figure S1. xmsPANDA output of the Type 1 Manhattan Plot (VIP vs mz).

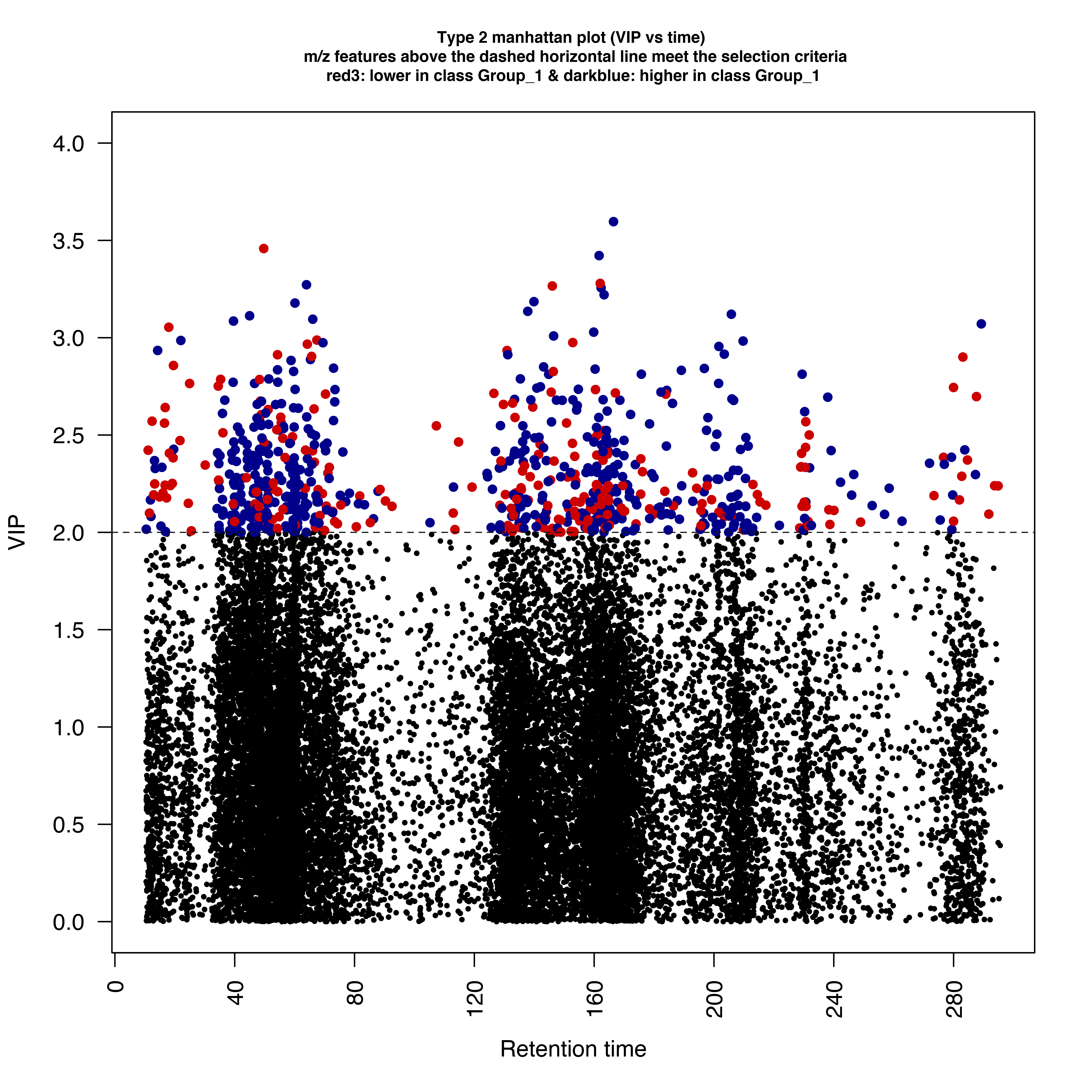

Figure S2. xmsPANDA output of the Type 2 Manhattan Plot (VIP vs time).

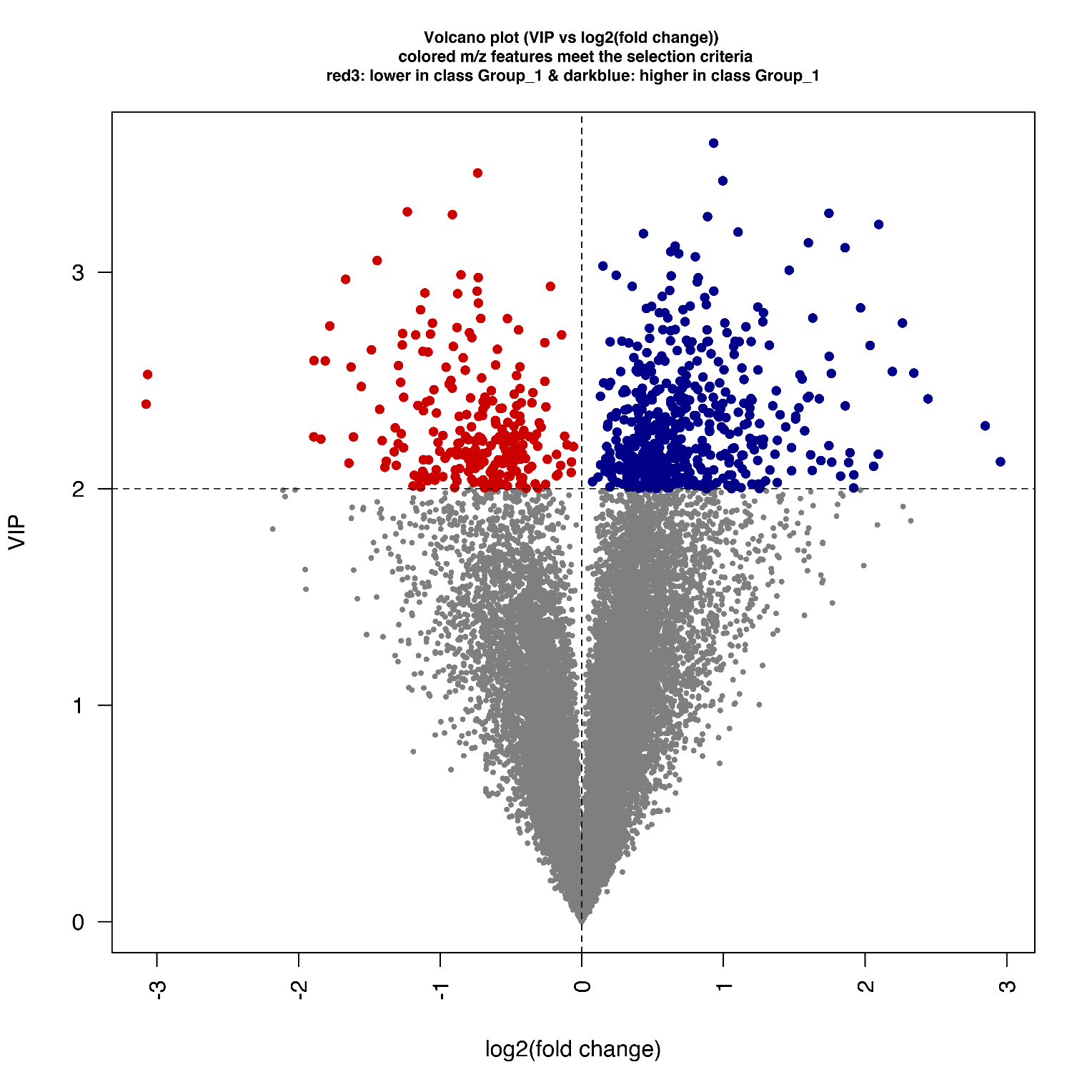

Figure S3. xmsPANDA output Volcano Plot (VIP vs log2FC).

| Table S1: Annotated Metabolites of Interest | | | | |
| --- | --- | --- | --- | --- |
| Query | Match | HMDB | PubChem | KEGG |
| C05645 | 4-(2-Amino-3-hydroxyphenyl)-2,4-dioxobutanoic acid | HMDB0004083 | 440741 | C05645 |
| C01252 | 4-(2-Aminophenyl)-2,4-dioxobutanoic acid | HMDB0000978 | 472 | C01252 |
| C14801 | 1-Nitro-5,6-dihydroxy-dihydronaphthalene | HMDB0060328 | 11954052 | C14801 |
| C00078 | L-Tryptophan | HMDB0000929 | 6305 | C00078 |
| C01345 | 2'-Deoxyinosine triphosphate | HMDB0003537 | 146302 | C01345 |
| C05598 | Phenylacetylglycine | HMDB0000821 | 68144 | C05598 |
| C16595 | 4-Hydroxy-5-phenyltetrahydro-1,3-oxazin-2-one | HMDB0060389 | 0 | C16595 |
| C16587 | 3-Carbamoyl-2-phenylpropionaldehyde | HMDB0060366 | 0 | C16587 |
| C01007 | Riboflavin reduced | HMDB0001557 | 22833571 | C01007 |
| C05651 | 5-Hydroxykynurenine | HMDB0012819 | 440745 | C05651 |
| C03227 | L-3-Hydroxykynurenine | HMDB0011631 | 11811 | C03227 |
| C16677 | 4-Hydroxyretinoic acid | HMDB0006254 | 6438629 | C16677 |
| C16679 | all-trans-18-Hydroxyretinoic acid | HMDB0012452 | 6506224 | C16679 |
| C16680 | all-trans-5,6-Epoxyretinoic acid | HMDB0012451 | 5363137 | C16680 |
| C00108 | 2-Aminobenzoic acid | HMDB0001123 | 227 | C00108 |
| C00461 | Chitin | HMDB0003362 | 21252321 | C00461 |
| C00294 | Inosine | HMDB0000195 | 6021 | C00294 |
| C00270 | N-Acetylneuraminate | NA | 3568 | C00270 |
| C01120 | Sphinganine 1-phosphate | HMDB0001383 | 644260 | C01120 |
| C02514 | 3-Fumarylpyruvate | HMDB0060371 | 5280525 | C02514 |
| C16741 | 5-Hydroxylysine | HMDB0000450 | 4433 | C16741 |
| C00780 | Serotonin | HMDB0000259 | 5202 | C00780 |
| C00015 | Uridine 5'-diphosphate | HMDB0000295 | 6031 | C00015 |
| G00092 | NA | NA | NA | NA |
| C00073 | L-Methionine | HMDB0000696 | 6137 | C00073 |
| C05446 | 27-Deoxy-5b-cyprinol | HMDB0001231 | 193321 | C05446 |
| C14781 | 15H-11,12-EETA | HMDB0005050 | 11954042 | C14781 |
| C14813 | 11H-14,15-EETA | HMDB0004693 | 11954058 | C14813 |
| C05966 | 15(S)-HPETE | HMDB0004244 | 5280893 | C05966 |
| C05356 | 5(S)-Hydroperoxyeicosatetraenoic acid | HMDB0001193 | 5280778 | C05356 |
| C02165 | Leukotriene B4 | HMDB0001085 | 5283128 | C02165 |
| C14823 | 8(S)-HPETE | HMDB0004699 | 9548880 | C14823 |
| C14812 | 12(R)-HPETE | HMDB0004692 | 9548885 | C14812 |
| C05965 | 12(S)-HPETE | HMDB0004243 | 5280892 | C05965 |
| C01134 | Pantetheine 4'-phosphate | HMDB0001416 | 987 | C01134 |
| C05512 | Deoxyinosine | HMDB0000071 | 65058 | C05512 |
| C00449 | Saccharopine | HMDB0000279 | 160556 | C00449 |
| C05640 | Cinnavalininate | HMDB0004078 | 114918 | C05640 |
| C11133 | Estrone glucuronide | HMDB0004483 | 115255 | C11133 |
| C11132 | 2-Methoxyestrone 3-glucuronide | HMDB0004482 | 22833612 | C11132 |
| C00268 | 4a-Carbinolamine tetrahydrobiopterin | HMDB0002215 | 1.35E+08 | C00268 |
| C04244 | 6-Lactoyltetrahydropterin | HMDB0002065 | 169508 | C04244 |
| C02953 | Dihydrobiopterin | HMDB0000038 | 252 | C02953 |
| C21640 | 6-(1'-Hydroxy-2'-oxopropyl)-tetrahydropterin | NA | 3.41E+08 | C21640 |
| C01284 | Inositol 1,3,4,5,6-pentakisphosphate | HMDB0003529 | NA | C01284 |
| C00473 | Retinol | NA | 3756 | C00473 |
| C16682 | 9-cis-Retinol | HMDB0006217 | 9947823 | C16682 |
| C00899 | 11-cis-Retinol | HMDB0006216 | 5280382 | C00899 |
| C00460 | Deoxyuridine triphosphate | HMDB0001191 | 65070 | C00460 |
| C16617 | 6-Mercaptopurine ribonucleoside triphosphate | HMDB0060411 | 3036942 | C16617 |
| C05841 | Nicotinate D-ribonucleoside | HMDB0006809 | 161234 | C05841 |
| C02670 | D-Glucurono-6,3-lactone | HMDB0006355 | 439782 | C02670 |
| C03289 | L-xylo-Hexulonolactone | METPA0381 | NA | C03289 |
| C17432 | all-trans-Decaprenyl diphosphate | HMDB0059616 | 5462210 | C17432 |
| C00253 | Nicotinic acid | HMDB0001488 | 938 | C00253 |
| C07446 | Isonicotinic acid | HMDB0060665 | 5922 | C07446 |
| C01137 | S-Adenosylmethioninamine | HMDB0000988 | 439415 | C01137 |
| C05490 | 11-Dehydrocorticosterone | HMDB0004029 | 13783449 | C05490 |
| C04677 | AICAR | HMDB0001517 | 65110 | C04677 |
| C00328 | L-Kynurenine | HMDB0000684 | 161166 | C00328 |
| C05647 | Formyl-5-hydroxykynurenamine | HMDB0012948 | 440743 | C05647 |
| C01346 | dUDP | HMDB0001000 | 145729 | C01346 |
| C05699 | Selenocystathionine | HMDB0006343 | 98223 | C05699 |
| C05452 | 3a,7a-Dihydroxy-5b-cholestane | HMDB0006893 | 3080603 | C05452 |
| C11134 | Testosterone glucuronide | HMDB0003193 | 108192 | C11134 |
| C00214 | Thymidine | HMDB0000273 | 5789 | C00214 |
| C19566 | 4-Hydroxy-4-(methylnitrosoamino)-1-(3-pyridinyl)-1-butanone | HMDB0062404 | 53297439 | C19566 |
| C19563 | 4-[(Hydroxymethyl)nitrosoamino]-1-(3-pyridinyl)-1-butanone | HMDB0062382 | 53297437 | C19563 |
| C00166 | Phenylpyruvic acid | HMDB0000205 | 997 | C00166 |
| C02763 | Enol-phenylpyruvate | HMDB0012225 | 641637 | C02763 |
| C03451 | S-Lactoylglutathione | HMDB0001066 | 440018 | C03451 |
| C06196 | dIMP | HMDB0006555 | 91531 | C06196 |
| C03263 | Coproporphyrinogen III | HMDB0001261 | 321 | C03263 |
| C05768 | Coproporphyrinogen I | HMDB0002158 | 68271 | C05768 |
| C05921 | Biotinyl-5'-AMP | HMDB0004220 | 440839 | C05921 |
| C00179 | Agmatine | HMDB0001432 | 199 | C00179 |
| C19691 | Farnesylcysteine | HMDB0011627 | 6438372 | C19691 |
| C20239 | 6-Carboxy-5,6,7,8-tetrahydropterin | HMDB0060410 | 12463319 | C20239 |
| C00099 | Beta-Alanine | HMDB0000056 | 239 | C00099 |
| C00041 | L-Alanine | HMDB0000161 | 5950 | C00041 |
| C00213 | Sarcosine | HMDB0000271 | 1088 | C00213 |
| C01888 | Aminoacetone | HMDB0002134 | 215 | C01888 |
| C05665 | 3-Aminopropionaldehyde | HMDB0001106 | 75 | C05665 |
| C01530 | Stearic acid | HMDB0000827 | 5281 | C01530 |
| C01801 | Deoxyribose | HMDB0003224 | 22833604 | C01801 |
| C00141 | Alpha-ketoisovaleric acid | HMDB0000019 | 49 | C00141 |
| C16360 | 3,7-Dimethyluric acid | HMDB0001982 | 83126 | C16360 |
| C16356 | 1,7-Dimethyluric acid | HMDB0011103 | 91611 | C16356 |
| C03150 | Nicotinamide riboside | HMDB0000855 | 439924 | C03150 |
| C03684 | Dyspropterin | HMDB0001195 | 128973 | C03684 |
| C00835 | Sepiapterin | HMDB0000238 | 65253 | C00835 |
| C04874 | 7,8-Dihydroneopterin | HMDB0002275 | 659 | C04874 |
| C00735 | Cortisol | HMDB0000063 | 657311 | C00735 |
| C01124 | 18-Hydroxycorticosterone | HMDB0000319 | 11222 | C01124 |
| C05469 | 17a,21-Dihydroxy-5b-pregnane-3,11,20-trione | HMDB0006758 | 65554 | C05469 |
| C06055 | O-Phospho-4-hydroxy-L-threonine | HMDB0006802 | 440901 | C06055 |
| C14857 | 1,1-Dichloroethylene epoxide | HMDB0060333 | 119521 | C14857 |
| C14858 | 2,2-Dichloroacetaldehyde | HMDB0060357 | 6576 | C14858 |
| C14859 | Chloroacetyl chloride | HMDB0060452 | 6577 | C14859 |
| C05335 | Selenomethionine | HMDB0003966 | 15103 | C05335 |
| C05673 | CMPciliatine | HMDB0060067 | 440754 | C05673 |
| C04494 | Guanosine 3'-diphosphate 5'-triphosphate | HMDB0060480 | 38166 | C04494 |
| C00438 | Ureidosuccinic acid | HMDB0000828 | 93072 | C00438 |
| C16241 | (R)-lipoic acid | HMDB0001451 | 6112 | C16241 |
| C00112 | CDP | HMDB0001546 | 6132 | C00112 |
| C00831 | Pantetheine | HMDB0003426 | 479 | C00831 |
| C00127 | Oxidized glutathione | HMDB0003337 | 975 | C00127 |
| C00122 | Fumaric acid | HMDB0000134 | 444972 | C00122 |
| C00500 | Biliverdin | HMDB0001008 | 5353439 | C00500 |
| C03492 | D-4'-Phosphopantothenate | HMDB0001016 | 131 | C03492 |
| C00319 | Sphingosine | HMDB0000252 | 5353955 | C00319 |
| C02934 | 3-Dehydrosphinganine | HMDB0001480 | 439853 | C02934 |
| C06429 | Docosahexaenoic acid | HMDB0002183 | 445580 | C06429 |
| C00881 | Deoxycytidine | HMDB0000014 | 13711 | C00881 |
| C14854 | 9-Hydroxybenzo[a]pyrene-4,5-oxide | HMDB0062439 | 115064 | C14854 |
| C02679 | Dodecanoic acid | HMDB0000638 | 3893 | C02679 |

Table S1: Annotated metabolites of interest.

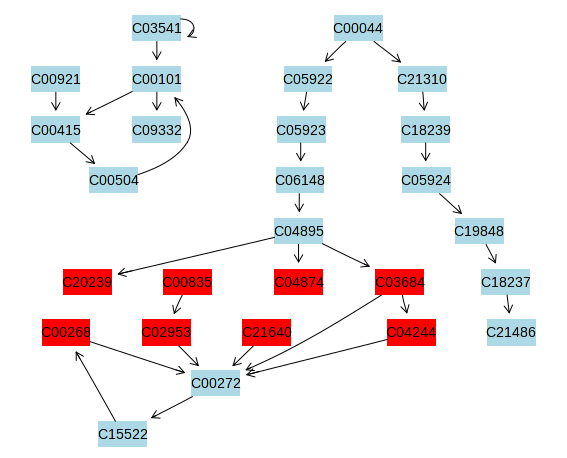

Figure S4. Folate Biosynthesis

| C03541 | THF-polyglutamate |
| --- | --- |
| C00101 | Tetrahydrofolate; 5,6,7,8-Tetrahydrofolate; Tetrahydrofolic acid; THF; (6S)-Tetrahydrofolate; (6S)-Tetrahydrofolic acid; (6S)-THFA |
| C09332 | THF-L-glutamate |
| C00504 | Folate |
| C00921 | Dihydropteroate; 7,8-Dihydropteroate |
| C00415 | Dihydrofolate; Dihydrofolic acid; 7,8-Dihydrofolate; 7,8-Dihydrofolic acid; 7,8-Dihydropteroylglutamate |
| C00044 | GTP |
| C05922 | Formamidopyrimidine nucleoside triphosphate |
| C05923 | 2,5-Diaminopyrimidine nucleoside trisphosphate |
| C06148 | 2,5-Diamino-6-(5’-triphosphoryl-3’,4’-trihydroxy-2’-oxopentyl)-amino-4-oxopyrimidine |
| C04895 | 7,8-Dihydroneopterin 3’-triphosphate; 2-Amino-4-hydroxy-6-(erythron-1,2,3-trihydroxypropyl) dihydropteridine triphosphate; 6-(L-erythro-1,2-Dihydroxypropyl 3-triphosphate)-7,8-dihydropterin; 6-[(1S,2R)-1-2-Dihydroxy-3-triphosphooxypropyl]-7,8-dihydropterin |
| C21310 | (8S)-3’,8-Cyclo-7,8-dihydroguanosine 5’-triphosphate |
| C18239 | Precursor Z |
| C05924 | Molybdopterin |
| C19848 | Adenylated molybdopterin |
| C18237 | Molybdoenzyme molybdenum cofactor |
| C21486 | Thio-molybdenum cofactor |
| C20239 | 6-Carboxy-5,6,7,8-tetrahydropterin; 6-Carboxytetrahydropterin |
| C00835 | Sepiapterin |
| C04874 | 7,8-Dihydroneopterin; Dihydroneopterin; 2-Amino-4-hydroxy-6-(D-erythro-1,2,3-trihydroxypropyl)-7,8-dihydropteridine |
| C03684 | 6-Pyruvoyltetrahydropterin; 6-(1,2-Dioxopropyl)-5,6,7,8-tetrahydropterin; 6-Pyruvoyl-5,6,7,8-tetrahydropterin |
| C00268 | Dihydrobiopterin; 6,7-Dihydrobiopterin; Quinoid-dihydrobiopterin; (6R)-6-(L-erythro-1,2-Dihydroxypropyl)-7,8-dihydro-6H-pterin |
| C02953 | 7,8-Dihydrobiopterin; L-erythro-7,8-Dihydrobiopterin |
| C21640 | 6-(1’-Hydroxy-2’-oxopropyl)-tetrahydropterin |
| C04244 | 6-Lactoyl-5,6,7,8-tetrahydropterin |
| C00272 | Tetrahydrobiopterin; 5,6,7,8-Tetrahydrobiopterin; 2-Amino-6-(1,2-dihydroxypropyl)-5,6,7,8-tetrahydoro-4(1H)-pteridinone; L-erythro-Tetrahydrobiopterin |
| C15522 | 4a-Hydroxytetrahydrobiopterin; 4a-Hydroxy-5,6,4,8-tetrahydrobiopterin; (6R)-6-(L-erythro-1,2-Dihydroxypropyl)-5,6,7,8-tetrahydro-4a-hydroxypterin |

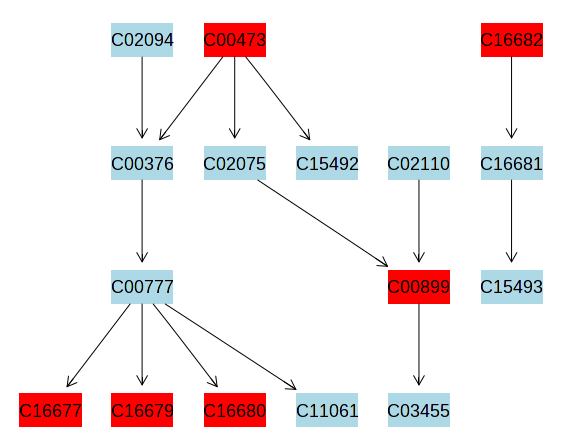

Figure S5. Retinol Metabolism

| C02094 | Beta-Carotene |
| --- | --- |
| C00473 | Retinol |
| C16682 | 9-cis-Retinol |
| C00376 | Retinal |
| C02075 | Retinyl ester |
| C15492 | All-trans-13,14-Dihydroretinol; 13,14-Dihydroretinol |
| C02110 | 11-cis-Retinal |
| C16681 | 9-cis-Retinal |
| C00777 | Retinoate; Retinoic acid; Vitamin A acid; all-trans-Retinoate; Acide retinoique (French) (DSL); Tretinoine (French) (EINECS); 3,7-Dimethyl-9-(2,6,6-trimethyl-1-cyclohexene-1-yl)-2,4,6,8-nonatetraenoic acid (ECL); (all-E)-3,7-Dimethyl-9-(2,6,6-trimethyl-1-cyclohexen-1-yl)-2,4,6,8-nonatetraenoic acid; beta-Retinoic acid; AGN 100335; all-(E)-Retinoic acid; all-trans-beta-Retinoic acid; all-trans-Retinoic acid; all-trans-Tretinoin; all-trans-Vitamin A acid; Ro 1-5488; trans-Retinoic acid; Tretin M; all-trans-Vitamin A1 acid |
| C00899 | 11-cis-Retinol |
| C15493 | 9-cis-Retinoic acid |
| C16677 | All-trans-4-Hydroxyretinoic acid |
| C16679 | all-trans-18-Hydroxyretinoic acid |
| C16680 | all-trans-5,6-Epoxyretinoic acid; all-trans-5,6-Epoxy-5,6-dihydroretinoic acid |
| C11061 | all-trans-Retinoyl-beta-glucuronide |
|  | 11-cis-Retinyl palmitate |

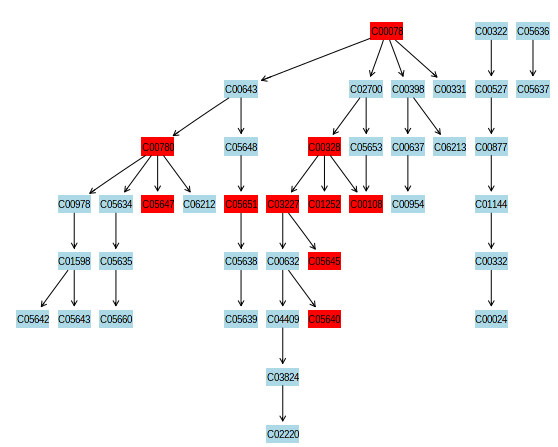

Figure S6. Tryptophan Metabolism

| C00078 | L-Tryptophan |
| --- | --- |
| C00322 | 2-Oxoadipate |
| C05636 | 3-Hydroxykynurenamine |
| C00643 | 5-Hydroxy-L-tryptophan |
| C02700 | L-Formylkynurenine |
| C00398 | Tryptamine |
| C00331 | Indolepyruvate |
| C00527 | Glutaryl-CoA |
| C05637 | 4,8-Dihydroxyquinoline; Quinoline-4,8-diol |
| C00780 | Serotonin |
| C05648 | 5-Hydroxy-N-formylkynurenine |
| C00328 | L-Kynurenine |
| C05653 | Formylanthranilate |
| C00637 | Indole-3-acetaldehyde |
| C06213 | N-Methyltryptamine |
| C00877 | Crotonoyl-CoA |
| C00978 | N-Acetylserotonin |
| C05634 | 5-Hydroxyindoleacetaldehyde |
| C05647 | Formyl-5-hydroxykynurenamine |
| C06212 | N-Methylserotonin |
| C05651 | 5-Hydroxykynurenine |
| C03227 | 3-Hydroxy-L-kynurenine |
| C01252 | 4-(2-Aminophenyl)-2,4-dioxobutanoate |
| C00108 | Anthranilate |
| C00954 | Indole-3-acetate |
| C01144 | (S)-3-Hydroxybutanoyl-CoA |
| C01598 | Melatonin |
| C05635 | 5-Hydroxyindoleacetate |
| C05638 | 5-Hydroxykynurenamine |
| C00632 | 3-Hydroxyanthranilate |
| C05645 | 4-(2-Amino-3-hydroxyphenyl)-2,4-dioxobutanoate |
| C00332 | Acetoacetyl-CoA |
| C05642 | Formyl-N-acetyl-5-methoxykynurenamine |
| C05643 | 6-Hydroxymelatonin |
| C05660 | 5-Methoxyindoleacetate |
| C05639 | 4,6-Dihydroxyquinoline; Quinoline-4,6-diol |
| C04409 | 2-Amino-3-carboxymuconate semialdehyde |
| C05640 | Cinnavalininate |
| C00024 | Acetyl-CoA |
| C03824 | 2-Aminomuconate semialdehyde |
| C02220 | 2-Aminomuconate |
